# Polyamines support myogenesis by facilitating myoblast migration

**DOI:** 10.1101/280206

**Authors:** Shirley Brenner, Yulia Feiler, Chaim Kahana

**Affiliations:** Department of Molecular Genetics, the Weizmann Institute of Science, Rehovot 76100, Israel

**Keywords:** Polyamines, differentiation, myoblast, migration, Ornithine decarboxylase, DFMO, myogenesis, HGF, Annexin A1

## Abstract

The regeneration of the muscle tissue relies on the differentiation of myoblasts into myocytes, to create myotubes and myofibers. Disruption of key events in this process may interfere with the correct formation or repair of muscle tissue. Polyamines, ubiquitous polycations that are essential for fundamental cellular processes, were demonstrated necessary for myogenesis; however, the mechanism by which polyamines contribute to this process has not yet been deciphered. In the present study, we examined the effect of polyamine depletion on the muscle regeneration model of C_2_C_12_ myoblasts. Our results reveal a requirement for polyamines at the very beginning of the muscle differentiation process. Myogenesis is accompanied by polyamine synthesis, even though the myoblasts contain high levels of polyamines at the moment of induction. Polyamine depletion at the time of induction, or inability to synthesize more polyamines during the first 24 hours of the process, inhibited myogenesis. Polyamine depletion inhibited the expression of all tested myogenic markers (Pax7, MyoD, Myogenin, Myf5 and Myosin heavy chain), as well as the cells migration and fusion abilities. Real time PCR analysis revealed two key early activation and migration factors, HGF and Annexin A1.

## INTRODUCTION

The adult skeletal muscle tissue has extensive intrinsic regeneration abilities. Regenerative myogenesis relies on the differentiation of myoblasts into myocytes, which fuse together to create myotubes and myofibers. Muscle stem cells are required to respond rapidly to injury. Upon exercise or when injury occurs, progenitor satellite cells will exit their normal quiescent state and start proliferating. Then, they transform to myoblasts, stop cycling, migrate, adhere and fuse to create multi-nuclei myotubes (1, 2). Several degenerative conditions affect the functionality of satellite cells, and thus harm the progression of muscle regeneration. At the molecular level, muscle stem cells, which express the transcription factors Pax3 and Pax7, are specified to form quiescent muscle satellite cells. These cells are maintained in a niche, shielded from most extracellular stimuli, and are activated only under specific conditions such as overwork, trauma or disease. The identity of the factors which activate satellite cells is gradually being deciphered, amongst the few known are hepatocyte growth factor (HGF) and nitric oxide (NO) (3–5). When activated, muscle satellite cells proliferate and co-express Pax7 and MyoD. Then, the majority of the population will commit to the myoblast lineage by down regulating Pax7, and up regulating Myf5 and MyoD, the main transcriptional activators of the myogenic regulatory factor family (MRF). Myf5 and/or MyoD are essential for skeletal muscle formation and regeneration (6–9). Later in the differentiation process myocytes must express the late MRFs Myogenin and MRF4, followed by the expression of muscle specific genes such as Myosin heavy chain (MHC) and muscle creatine kinase (MCK), which serve as markers for the completion of the differentiation process (2, 6, 10–16).

C_2_C_12_ mouse myoblasts, an established cell line derived from muscle satellite cells, serves as a model for studying myogenic differentiation, since they can be rapidly and efficiently induced to differentiate into multinucleated myotubes and myofibers under condition of serum deprivation (17–19)

The polyamines spermidine and spermine and their diamine precursor, putrescine, are ubiquitous polycations that are essential for fundamental cellular processes, such as transcription, translation and cellular proliferation, and differentiation (20). Putrescine is formed by the action of ornithine decarboxylase (ODC), the first and rate limiting enzyme in the polyamine biosynthesis pathway. This highly regulated enzyme is therefore critical for maintaining optimal polyamine levels (21, 22). Treatment with the irreversible, mechanism based ODC inhibitor, c (DFMO), results in polyamine depletion and inhibition of cellular proliferation and differentiation (23–27). Past studies implicated a role for polyamines in myogenesis (28), however, the ways polyamines participate in this differentiation process remain mostly obscure.

In the present study we investigated the effect of polyamine depletion on the differentiation of C_2_C_12_ myoblasts. We demonstrate here that polyamines are essential for muscle regeneration, acting at the onset of this process. Polyamine depletion inhibited the formation of myotubes and myofibers, distorting the correct expression pattern of myogenesis regulators and executers throughout the process. Furthermore, we show that polyamine depletion also interferes with cell migration, probably due to obstructed expression of key activation and migration regulators.

## RESULTS

### Polyamines are required for C_2_C_12_ cells differentiation

To determine how polyamine depletion affects myogenic differentiation, DFMO was added to the growth medium of C_2_C_12_ cells two days prior to their induction. The cells were induced to differentiate at 70-80% confluence using serum free differentiation medium (DM) containing or lacking DFMO. Differentiation was assessed 5 days post induction. While efficient differentiation was clearly observed in control cells, the differentiation was completely inhibited by the DFMO treatment (Fig 1A). To confirm that DFMO inhibited cellular differentiation solely through polyamine depletion, spermidine was added to the DM together with DFMO. As shown in figure 1, the inhibitory effect of DFMO was fully reversed by the addition of spermidine (10µM). Following stimulation, DFMO treated myoblasts failed to express the typical changes observed in control cells: the down regulation of Pax7 was delayed, MyoD and Myf5 failed to accumulate, and Myogenin was expressed to a much lower level compared to DFMO untreated cells. Lastly, there was almost complete elimination of MHC, the marker of terminally differentiated myocytes. Supplemented spermidine restored the manifestation of these changes (Fig 1B,C). Interestingly, to some extent the added spermidine led to a more profound expression of the myogenic markers: Pax7 was downregulated earlier, and Myogenin and MHC were expressed earlier and to higher levels (Fig 1B, C).

**Figure 1:**
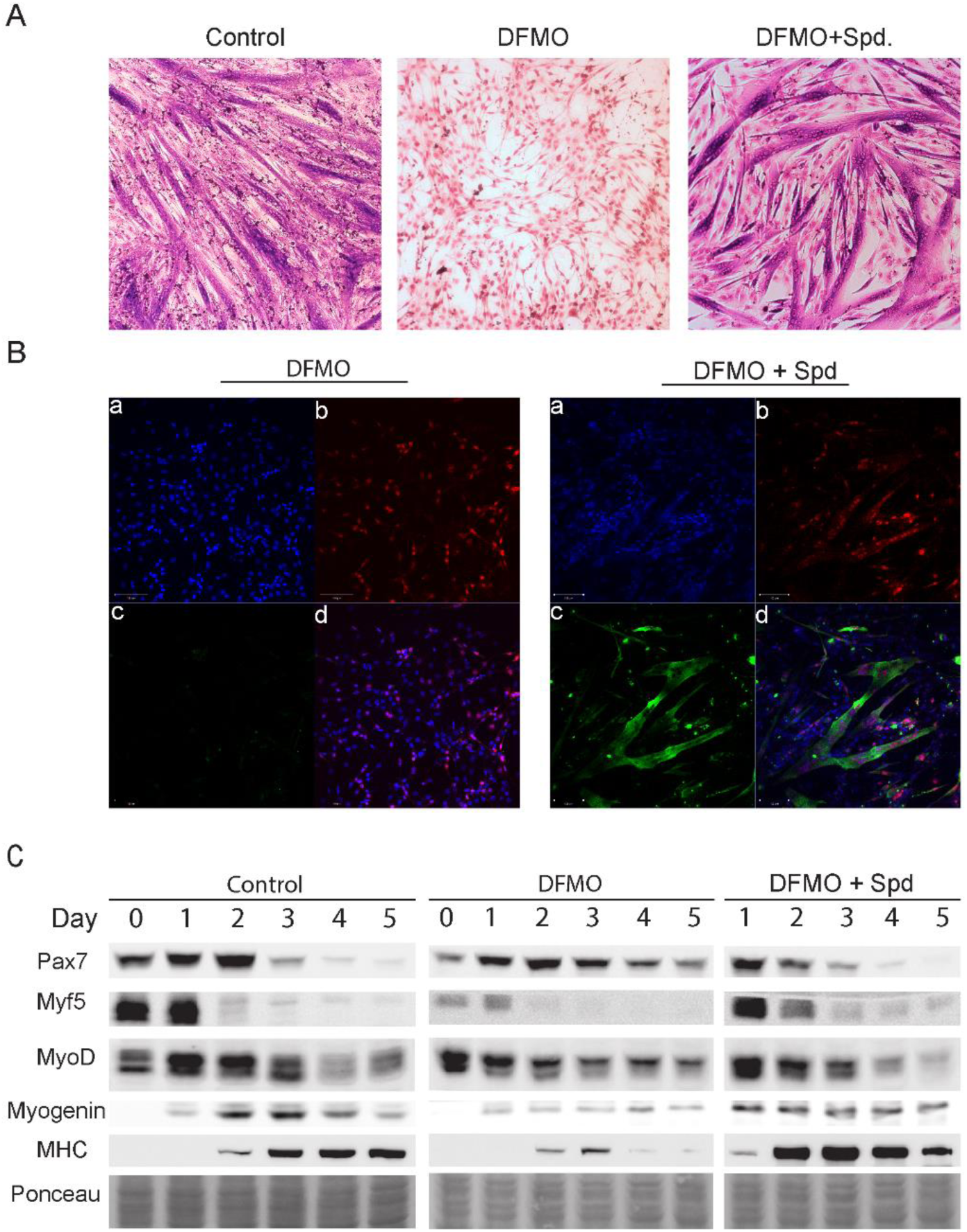
Polyamines are required for differentiation of C_2_C_12_ myoblasts. C_2_C_12_ myoblasts were induced to differentiate for 5 days with DM (Control), in the presence of DFMO (5mM), or DFMO (5mM) + spermidine (10μM), as indicated. (A) The cells were stained using the May-Grünwald – Giemsa method, as described under “Experimental Procedures”. Protein reach myotubes are identified by a dark purple color, while myoblasts (nuclei) are lightly stained in pink. (B) C_2_C_12_ myoblasts were grown on slides, fixed after 5 days of incubation with differentiation medium, and immunostained by (b) Mouse α myogenin and (c) Mouse α MHC antibodies, followed by staining with Cy5-Goat α Mouse (red) and Cy3-Goat α Mouse (green), respectively. (a) The cells were then mounted by ProLong (r) Gold antifade reagent, containing DAPI (Blue). (d) Merge. (C) Cellular extracts were prepared at the indicated times. The levels of individual proteins were determined by Western blot analysis.

### ODC activity and polyamines are induced by myogenic stimulation

To further investigate the requirement of polyamines in the myogenic differentiation process, we have determined ODC activity and polyamine levels along the standard myogenic induction protocol. C_2_C_12_ myoblasts were stimulated at 70-80% confluence and ODC activity and polyamines were determined in cellular extracts prepared at the indicated times following stimulation. A single sharp pick of ODC activity was observed one day after induction, and declined thereafter. This increase in ODC activity was completely abolished in the DFMO pretreated cells (Fig 2A). In control cells the level of polyamines was high already at the beginning of the process, slightly increased at the first day post stimulation (paralleling the rise in ODC activity) and declined thereafter. In the DFMO pretreated cells the polyamines (putrescine and spermidine but not spermine) were depleted already at the beginning of the process and their level was not increased at day 1 of the differentiation process (Fig 2B).

**Figure 2:**
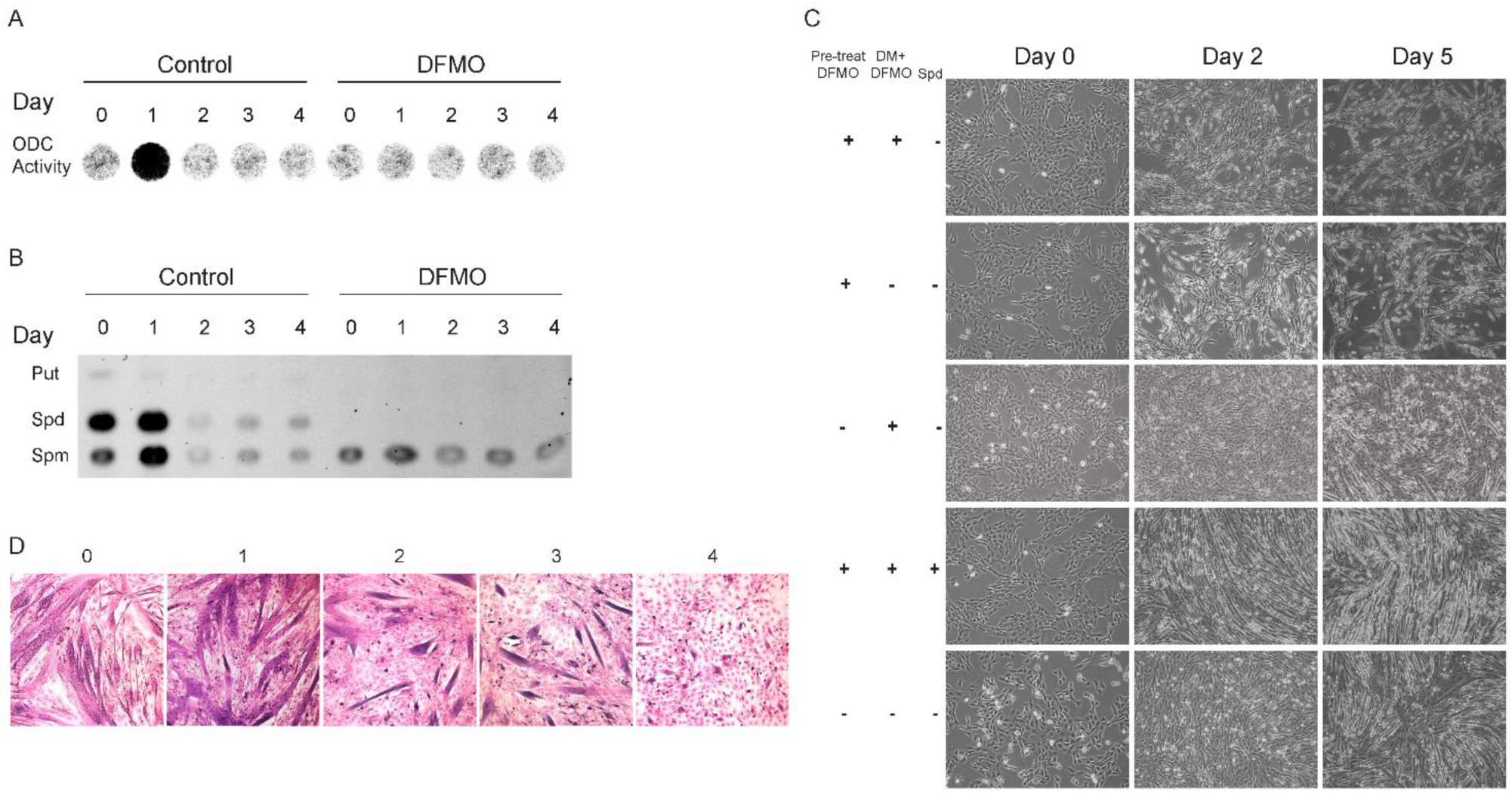
ODC activity and polyamines are required at the onset of myogenesis. C_2_C_12_ myoblasts were induced to differentiate with DM (Control), in the presence or absence of DFMO (5mM) and/or spermidine (10μM), as indicated. (A, B) DFMO (5mM) was added to the cells two days before induction and cellular extracts were prepared at the indicated times. Cellular extracts were prepared and subjected to ODC activity assay (A) and polyamine analysis (B) as described under “Experimental Procedures”. The positions of polyamine markers are indicated as *Put*, putrescine; *Spd*, spermidine; *Spm*, spermine. (C) C_2_C_12_ cells were treated as indicated on the left. The cultures were photographed at the indicated times, using Olympus DP73 camera adapted to Olympus IX73 microscope. *Pre-treat DFMO*, DFMO was added to the culture two days prior to induction; *DM+ DFMO*, DFMO was added to the differentiation medium at day 0 onward; *Spd*, Spermidine was added to the differentiation medium at day 0 onward. (D) C_2_C_12_ myoblasts were induced to differentiate in the presence of DFMO. Spermidine was added to the medium at the indicated times. The cells were stained using the May-Grünwald – Giemsa method, as described under “Experimental Procedures”.

While pre-treatment with DFMO inhibited the myogenic differentiation, addition of Spermidine together with DFMO at the day of induction restored the manifestation of successful differentiation. Stimulation of polyamine depleted (2 days DFMO pretreated) cells with DM lacking DFMO failed to induce differentiation, suggesting that either the pre-existing polyamines or those induced at the first day post induction are required for the outset of myogenesis (Fig 2C, second panel). In order to determine whether the increase in ODC activity and polyamines observed one day post induction is required for the differentiation process, DFMO was added four hours prior to the differentiation stimulus. Although some cell fusions were observed, the cells were unable to create myotubes and fully differentiate (Fig 2C, third panel), suggesting that the polyamines synthesized at the first day following induction are required for the establishment of efficient differentiation. In agreement with what we observed molecularly (see Fig 1C), addition of spermidine to the differentiation medium together with DFMO resulted in faster differentiation, i.e. some cell fusions were already observed at day 2 post induction (Fig 2C, fourth panel). To gain better insight regarding the stage of the differentiation process at which polyamines are required, we added spermidine on consecutive days following myoblasts stimulation. Addition of spermidine at days 0 and 1 overcame the inhibitory effect of DFMO, but when added at day 2 onwards, it was practically ineffective (Fig 2D). Taken together, these results indicate that polyamines are required at the very beginning of the myogenic process.

### Polyamines are required for the differentiation of confluent C_2_C_12_ culture

C_2_C_12_ keep dividing after they are induced, until they exit the cell cycle and fuse (1) (see fig 2C, bottom row). Therefore, we considered the possibility that polyamines are required in this differentiation process through their requirement for cellular proliferation. The proliferative step may be directly required for the myogenic differentiation as demonstrated in adipogenesis (26, 29), or it may be needed to establish a tight contact between the cells in order to facilitate their fusion. To examine these possibilities, C_2_C_12_ myoblasts were grown to confluence and then induced to differentiate. In agreement with Tanaka et. Al (30), the confluent culture differentiated very efficiently, and to some extent exhibited a more rapid expression pattern of the myogenesis markers (Fig 3A compared to Fig 1C). Also in the confluent cultures, an increase in ODC activity and polyamine level was observed one day post induction. DFMO completely inhibited ODC activity, and depleted cellular polyamines similarly to what was observed in the stimulated sub confluent culture (Fig 3B, C compared to Fig 2A, B). To determine if cell proliferation is required for the induction of myogenesis in confluent culture, we conducted a ^3^H-thymidine incorporation assay following the myogenic stimulation. For this purpose, the cells were labeled with ^3^H-thymidine for 40 hours following their induction. As can be seen in figure 3D, there were no cell divisions following myogenic stimulation of the confluent cultures. Polyamine depleted C_2_C_12_ myoblasts were unable to differentiate even when a confluent culture was induced, and this inhibition was reversed by the addition of spermidine to the DM (Fig 3E). These results suggest that polyamine depletion obstructs myoblasts differentiation regardless of the polyamines effect on cell divisions, and that prevention of cell-cell contact cannot account for this inhibition.

**Figure 3:**
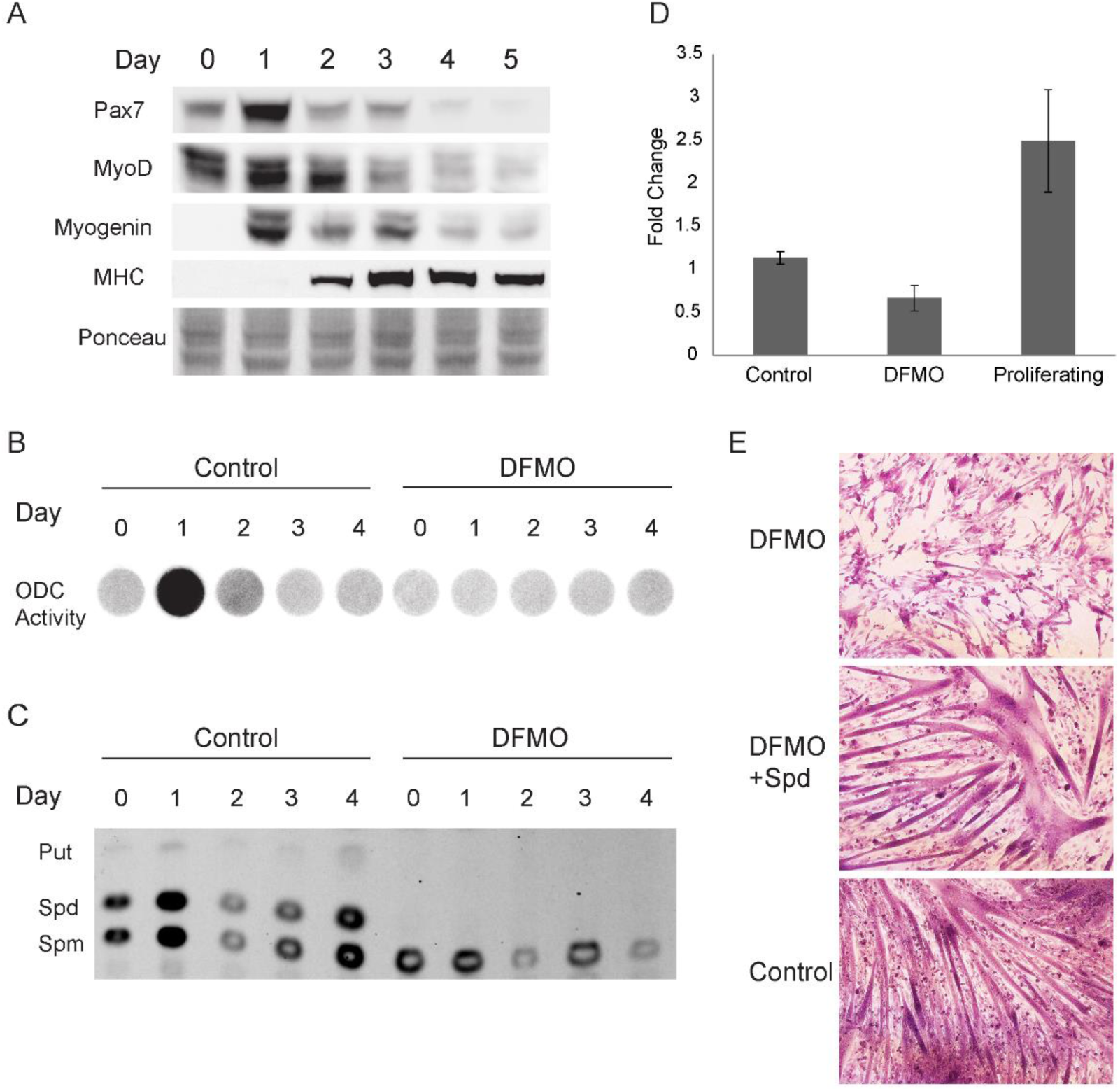
Polyamines are required for the differentiation of confluent C_2_C_12_ cultures. C_2_C_12_ myoblasts were grown to confluence and induced to differentiate with DM (Control), in the presence or absence of DFMO (5mM) and/or spermidine (10μM), as indicated. DFMO was added to the culture two days before induction. (A) Cellular extracts were prepared at the indicated times. The levels of individual proteins were determined by Western blot analysis. (B, C) DFMO (5mM) was added to the cells two days before induction, cellular extracts were prepared at the indicated times and subjected to ODC activity assay (B) and polyamine analysis (C) as described under “Experimental Procedures”. The positions of polyamine markers are indicated. *Put*, putrescine; *Spd*, spermidine; *Spm*, spermine. (D) Cells were labeled with H^3^-thymidine from the induction time for 40 hours. The presented histogram represents fold change between induced and non-induced cells. *Control*, C_2_C_12_ myoblasts induced with regular DM; *DFMO*, C_2_C_12_ myoblasts pretreated with DFMO were induced with DM + DFMO (5mM), *proliferating*, quiescent NIH-3T3 cells (maintained in serum poor (2%) medium), were induced to divide synchronously with serum rich (20%) medium. (E) The cells were stained using the May-Grünwald – Giemsa method, as described under “Experimental Procedures”. Each well was photographed randomly using Olympus DP73 camera adapted to Olympus IX73 microscope. Where indicated, DFMO was added to the culture two days before induction, spermidine was added at the day of induction.

### Polyamine depletion inhibits cell motility and cell fusion

During their differentiation process, myoblasts differentiate into myocytes, which fuse to form multinucleated myotubes and myofibers. In order for the fusion to occur, the cells send pseudopodia to contact each other, migrate, align, adhere and then fuse (2). The inability of polyamine depleted C_2_C_12_ myoblasts to differentiate, even when they are in tight contact, raises the possibility that these cells might not be able to perform any of the next required steps (i.e. migration, alignment, adherence and/or fusion). To investigate this hypothesis, live imaging was performed to follow the cells migratory behavior following their induction. Stimulated cells were subjected to time laps microscopy for 16 hours, starting 24 hours post induction. Under standard differentiation conditions, C_2_C_12_ myoblasts send pseudopodia, migrate, and some adhere and start fusing. In contrast, while polyamine depleted cells do send pseudopodia, they elongate to a thread-like shape and fail to migrate. Addition of spermidine to the DFMO treated cells enabled them to regain their rounded morphology, and restored their ability to migrate and fuse (Fig 4A, videos 1-3).

**Figure 4:**
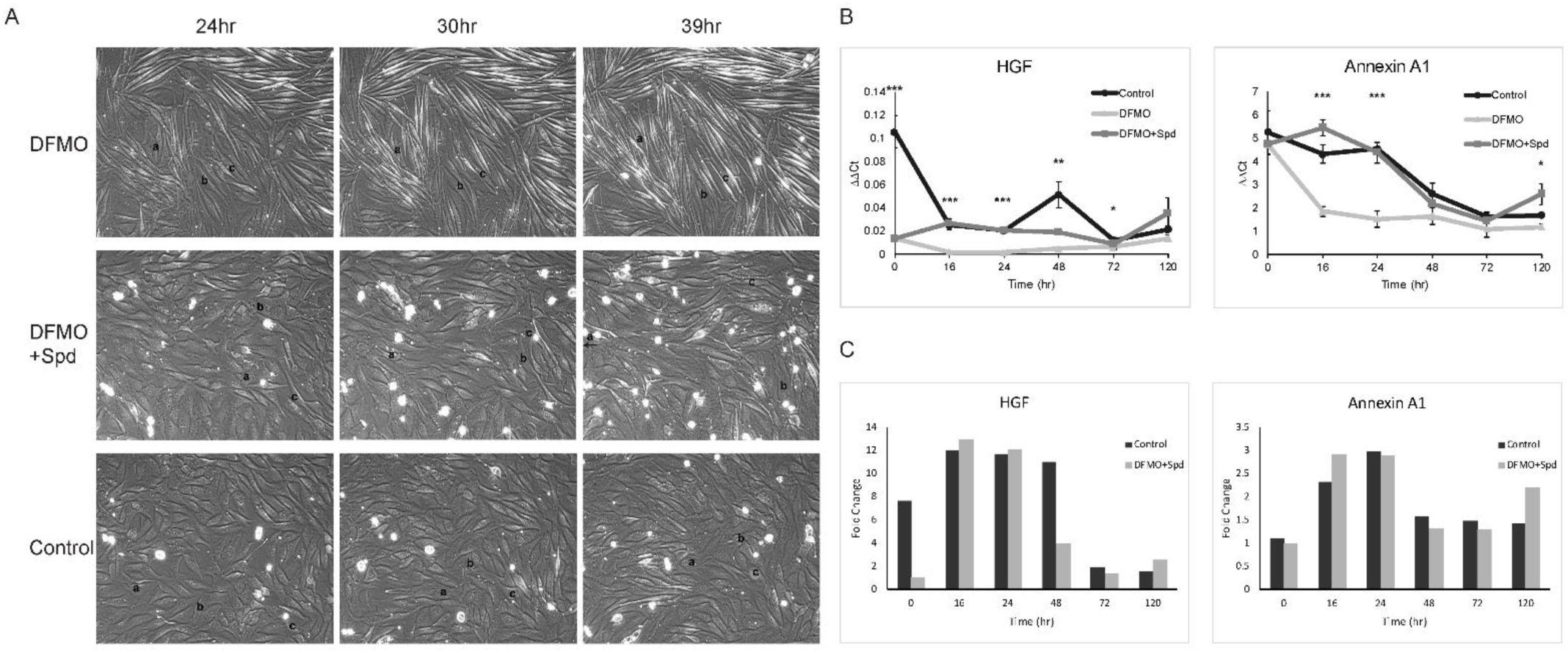
Polyamine depletion inhibits cell motility and cell fusion. C_2_C_12_ myoblasts were induced to differentiate with DM (Control), in the presence or absence of DFMO (5mM) and/or spermidine (10μM), as indicated. (A) The cells were subjected to time laps microscopy every 10 minutes for 16 hours, starting 24 hours post induction. Three individual photos from the indicated times are shown, in each panel the same three cells were marked (a, b, c, arrow marks a cell that went out of the frame). See full videos at the supplementary data. (B) Total RNA was isolated at the indicated times and mRNA levels were measured by quantitative real-time PCR. Data are presented as ∆∆Ct means of three independent experiments. Standard deviations are represented by error bars. Statistically significant mRNA levels of DFMO treated cells are marked with asterisks. (C) Histograms show ∆∆Ct fold Change versus DFMO treated cells.

Myoblasts activation and migration depend on the expression of a number of factors. We have employed real-time PCR to determine the level of several mRNAs encoding proteins relevant to the process of cellular migration and myogenesis. As shown in figure 4B, the level of hepatocyte growth factor (HGF), which is a known activator of satellite cells and myoblasts migration (31–34), was about 7 fold lower in DFMO treated cells compared to control cells on the day of induction. At 16 hours, the mRNA level of HGF in control cells was down regulated, but was still about 12 fold higher than its level in DFMO treated cells, and remained about 11 fold higher throughout the first 48 hours of the differentiation process. Addition of Spermidine to the DFMO treated cells led to recovery of HGF mRNA levels, reaching more than 12 fold already at 16 hours post induction. While similar levels of Annexin A1mRNA, a factor necessary for myoblasts migration (35, 36) were monitored in control and polyamine depleted myoblasts at time 0, polyamine depletion led to down regulation of Annexin A1 mRNA level by 2-3 fold at 16-24 hours post induction (Fig 4B, C). Addition of spermidine to the DM led to complete recovery of Annexin A1 mRNA levels, reaching about 3 fold higher level already 16 hours post induction (Fig 4B, C). Other genes that were examined (e.g. IGFBP4, TnnT1, Bin3, Myostatin, RhoC, and RhoA) did not show any significant difference between control and DFMO treated cells. These results indicate that polyamines positively regulate mRNA levels of these two migration genes at the onset of myogenic differentiation.

## DISCUSSION

In the present study, we show that polyamine depletion inhibits muscle regeneration at its onset, by impairing the ability of the cells to migrate and fuse. Polyamines are necessary for proliferation and differentiation of mammalian cells (20, 27, 37–40). Here we demonstrate that myogenic regeneration is accompanied by the induction of ODC and de-novo synthesis of polyamines. Polyamine depletion halted C_2_C_12_ migration and differentiation, paralleled by perturbation of the expression of proteins required for the proper manifestation of this process.

C_2_C_12_ myoblasts have high levels of polyamines at the onset of myogenesis, and when induced they produce more polyamines due to elevation of ODC activity.

Polyamine depletion seems to inhibit myogenesis at the onset of the differentiation process. The level of Pax7 is low at the beginning, and its essential down regulation is impaired. This distorted expression pattern is followed by failure to up regulate MyoD, Myf5 and Myogenin. Addition of spermidine to the DFMO treated cells reversed this disruption, leading to a sharper differentiation pattern, i.e. faster down regulation of Pax7, and earlier and stronger expression of MHC. Spermidine can reverse the effect of DFMO only when added at the very beginning of the process, in agreement with early induction of ODC activity and polyamines during the differentiation process.

Under the standard myogenesis protocol some cellular proliferation still occurs during the first 24-48 hours post induction. This proliferation is inhibited by treatment with DFMO, and is restored by the addition of spermidine. Since after induction, C_2_C_12_ cells continue to proliferate before they exit the cell cycle and differentiate (1), and since polyamines were implicated in supporting cellular proliferation (27, 37), we considered the possibility that polyamines are required in myogenesis through their proliferative attribution. However, since two-day post confluent myoblasts were efficiently differentiated without demonstrating DNA synthesis, and since this differentiation was prevented by DFMO and restored by the addition of spermidine, we concluded that polyamines are required in this process not through supporting proliferation. The inhibition of myogenesis by DFMO in fully confluent C_2_C_12_ culture also suggests that polyamines are not required for generating cell-cell contact via their support of cellular proliferation.

Stimulation of polyamine depleted cells with medium lacking DFMO failed to induce differentiation. Induction of untreated cells with DM containing DFMO (i.e. the cells had normal levels of polyamines at the starting point but were unable to keep synthesizing them throughout the first 24 hours) also inhibited differentiation. We also demonstrated that polyamines reversed the effect of DFMO pretreatment only if added during the first day post induction. Together these results demonstrate that the process of myogenic differentiation requires polyamines synthesized following the myogenic stimulus, and that these polyamines act at the onset of this differentiation process.

In order to differentiate, myoblasts undergo sequential necessary steps: They migrate, align, send membrane extensions towards each other, adhere and then fuse to create multinucleated myotubes (2, 3, 15). Time lapse imaging of induced C_2_C_12_ cells revealed that polyamine depletion inhibits both their ability to migrate and their ability to fuse. While the DFMO treated cells elongate, they only gain a thread-like morphology, and unable to migrate or fuse. Addition of spermidine to the DFMO treated cells enabled the regaining of these critical steps. Examination of mRNA levels of motility related genes revealed that polyamines are necessary for the expression of HGF, which plays a role during migration and early patterning of muscle precursor cells leading to their commitment for the myogenic lineage, and stimulates migration of myoblasts (31–34) and Annexin A1, known to promote cell migration, and specifically promotes migration of myoblasts, hence positively regulating muscle cell differentiation (35, 36, 41). Moreover, these changes occur specifically at the beginning of the differentiation process, overlapping the time frame requirement for polyamines.

In conclusion, our study consistently demonstrates a need for polyamines at early stages of myogenesis. This is done by facilitating myoblasts migration during muscle regeneration, through positive regulation of migration related genes.

## EXPERIMENTAL PROCEDURES

### Cell culture conditions and induction of differentiation

C_2_C_12_ mouse myoblasts were grown in Dulbecco’s modified Eagle’s medium (DMEM, Invitrogen), supplemented with 15% (v/v) fetal bovine serum (Biological Industries), 100 units/ml penicillin, and 100 g/ml streptomycin (Pen-Strep, Biological Industries) at 37 °C in 7% CO_2_. Induction of differentiation was done by switching to serum deprived DMEM containing 1μg/ml insulin (Homulin, Lyli) denoted as DM. The medium was refreshed every other day.

### May-Grünwald – Giemsa staining

The cells were washed (PBS), fixed and stained with 0.25% May-Grünwald in methanol for 1 minute, washed with tap water, stained with Giemsa 1/10 in tap water for 5 minutes and washed again with tap water. For each experiment all of the samples were stained at the same time. Protein reach myotubes are identified by a dark purple color, while myoblasts (nuclei) are lightly stained in pink. Each well was photographed randomly using Olympus DP73 camera adapted to Olympus IX73 microscope.

### Immunoblot analyses

Cells were induced to differentiate as described above, and cellular extracts were prepared at the indicated time points. Cellular extracts were prepared by lysing cells in RIPA buffer (50mM Tris-HCl pH 8, 150mM KCl, 1.0% NP-40 (IGEPAL), 0.5% sodium deoxycholate, 0.1% SDS). Equal portions of protein were resolved by electrophoresis in SDS-PAGE, electroblotted to a nitrocellulose membrane and incubated with the indicated antibodies followed by HRP-conjugated anti-IgG antibodies. The antibodies used were: mouse hybridoma mAb anti-myosin heavy chain (DHSB), mouse hybridoma mAb anti-Pax7 (DHSB), mouse mAb anti-myogenin (Santa Cruz), rabbit pAb anti-MyoD (Santa Cruz), rabbit pAb anti-MyF5 (Santa Cruz).

### ODC activity assay

Cells were washed with PBS, suspended in ODC activity buffer (25 mM Tris-HCl pH 7.5, 2.5 mM DTT, 200 μM PLP) and lysed by three cycles of freeze/thaw. Lysates were centrifuged, supernatant was collected and protein concentration was determined using the Bradford reagent. A portion of 300μg protein was added to final volume of 90μl ODC activity buffer in a 96 well microtiter plate. C^14^-Ornithin (0.1μCi; Perkin Elmer) in ODC activity buffer was added to each well. Whatman 3mm filter paper impregnated with saturated BaOH was used to cover the microtiter plate. The closed plate was incubated for 3 hours at 37°C. The filter was washed with acetone and dried. Liberated C^14^-CO_2_ that was trapped in the filter paper was monitored using Typhoon FLA7000 phosphor-imaging scanner.

### Polyamine Analysis

The cells were harvested, pelleted, and suspended in 100μl PBS. The cells were lysed in 3% perchloric acid, and the precipitated material was removed by centrifugation (5 min at 13,000 rpm). The supernatant was collected for polyamine analysis, and the pellet was used for normalization by DNA quantification (DNA was quantified by suspending the pellet in 400μl of 4% diphenylamine (Sigma) in acetic acid, 400μl of 10% perchloric acid, and 20μl of 1:500 acetaldehyde (Sigma), and by incubation for 16h at 30 °C followed by absorbance determination at 595nm). For polyamine analysis, 100μl of the perchloric acid supernatant was mixed with 200μl of 3mg/ml dansyl-chloride (in acetone). The mixture was incubated for 16h in the dark with 100μl saturated sodium carbonate. To neutralize residual dansyl-chloride, 16.67mg/ml L-proline solution was added and incubated for 1h at room temperature. Dansylated derivatives were extracted into toluene by centrifugation. Samples were spotted on Silica 60 F_254_ TLC plate (Merck), and the dansylated derivatives were resolved by TLC using ethyl acetate/cyclohexane (1:1.5) as a solvent, in the dark. The plate was visualized by UV.

### Thymidine-H^3^ Incorporation Assay

Cells were pulsed for 40hr from the time of induction. Then, the cells were lysed in 0.5ml of 0.5M KOH and chromosomal DNA was precipitated with 1ml TCA 20%, 30min on ice. The DNA was filtered through GF/C filters, washed with TCA 20% and then with ethanol 96%, dried and counted in 5ml scintillation fluid.

### Time lapse microscopy

The cells were photographed using Zeiss Axiocam 506 mono, adapted to Zeiss Observer.Z1 microscope. Time lapse photos were taken starting 24 hours post induction, every 10 minute, for 16 hours, and were composed into movies using ZEN2 pro program.

### Real Time PCR

RNA was extracted from cells at the indicated time points using TRI-reagent. cDNA was synthesized using the qScript kit (Quanta Biosciences) followed by DNAse treatment. Real time PCR analysis was performed using SYBR Green I (Quanta Biosciences) as a fluorescent dye, according to the manufacturer’s instructions. All experiments were carried out in triplicates, and results were normalized to HPRT RNA. Real time PCR primers were selected from the Harvard Primer Bank (Primer Bank IDs are as follows: HPRT-7305155a1; AnxA1-6754570a1; HFG-4165287a1; IGFBP4-6981086a1; Bin3-118130496c1; TnnT1-6755841a1; RhoA-31542143a1; RhoC-6680728a1; Myostatin-6754752a1). Data was analyzed using the 2^-ΔΔCt^ method.

## Acknowledgements

We would like to thank Dr. Kfir Umansky for advice and suggestions on this work.

## FOOTNOTES

 [Funding was provided by: Israel Academy of Science and Humanities [855/15]]
 [The abbreviations used are: ODC, Ornithine decarboxylase; DFMO, α-difluoromethylornithine; DM, differentiation medium; MHC, myosin heavy chain; HGF, hepatocyte growth factor]

